# Children with amblyaudia show less flexibility in auditory cortical entrainment to periodic non-speech sounds

**DOI:** 10.1101/2021.09.08.459520

**Authors:** Sara Momtaz, Deborah Moncrieff, Meredith A. Ray, Gavin M. Bidelman

**Affiliations:** School of Communication Sciences & Disorders, University of Memphis, Memphis, TN, USA; Division of Epidemiology, Biostatistics, and Environmental Health, School of Public Health, University of Memphis, Memphis, TN, USA; Institute for Intelligent Systems, University of Memphis, Memphis, TN, USA; University of Tennessee Health Sciences Center, Department of Anatomy and Neurobiology, Memphis, TN, USA

**Keywords:** Auditory processing disorders (APD), event-related brain potentials (ERPs), gamma/beta band response, hemispheric asymmetries, phase-locking, time-frequency analysis

## Abstract

**Objective:** We investigated auditory temporal processing in children with amblyaudia (AMB), a subtype of auditory processing disorder, via cortical neural entrainment.

**Design and study samples:** Evoked responses were recorded to click-trains at slow vs. fast (8.5 vs. 14.9/sec) rates in n=14 children with AMB and n=11 age-matched controls. Source and time-frequency analyses decomposed EEGs into oscillations (reflecting neural entrainment) stemming from the bilateral auditory cortex.

**Results:** Phase-locking strength in AMB depended critically on the speed of auditory stimuli. In contrast to age-matched peers, AMB responses were largely insensitive to rate manipulations. This rate resistance was seen regardless of the ear of presentation and in both cortical hemispheres.

**Conclusion:** Children with AMB show a stark inflexibility in auditory cortical entrainment to rapid sounds. In addition to reduced capacity to integrate information between the ears, we identify more rigid tagging of external auditory stimuli. Our neurophysiological findings may account for certain temporal processing deficits commonly observed in AMB and related auditory processing disorders (APDs) behaviorally. More broadly, our findings may inform communication strategies and future rehabilitation programs; increasing the rate of stimuli above a normal (slow) speech rate is likely to make stimulus processing more challenging for individuals with AMB/APD.

## INTRODUCTION

Time representation and perception are fundamental cognitive skills of interest to both basic and clinical research. Temporal processing refers to the detection, identification, integration, and segregation of sound events over time (Picton, 2013). Temporal processing impairments are observed in a variety of patient populations such as schizophrenia (Luthra, 2021), Parkinson’s disease (Grondin, 2010), attention-deficit/hyperactivity disorder (ADHD) (Toplak et al., 2006), dyslexia (Tallal, 1980), language impairment (Dawes et al., 2009), and learning disorders (McFarland & Cacace, 2009). Difficulties “hearing in time” are particularly salient in children with auditory processing disorder (APD), where temporal processing deficits are reflected in poorer auditory perceptual abilities (Tallal, 1980; Merzenich et al., 1996; Picton, 2013). Presumably, temporal processing deficits during childhood negatively affect speech-language acquisition, which relies heavily on the accurate encoding of fine timing information in sound (Picton, 2013). This has led to the use of rhythm and time synchronization paradigms in rehabilitative and therapeutic approaches (Grondin, 2010) to improve both basic auditory and more general cognitive abilities such as attention and memory (Tallal, 1980; Picton, 2013). Nevertheless, due to the heterogeneity of APD, there might be a range of different temporal difficulties which span both speech and non-speech domains.

APD is characterized by symptoms of hearing difficulty without hearing loss (i.e., changes in peripheral sensitivity), *per se*. APD comprises 5% of clinical hearing referrals (Moore, 2006). Amblyaudia (AMB) is a subcategory of APD that is defined as an abnormally large asymmetry (>2 SD) in dichotic listening performance due to hypoactivity of the non-dominant ear or hyperactivity of the dominant ear (Moncrieff et al., 2016). Possible etiologies and underlying neural mechanisms in AMB are only beginning to be explored (Moncrieff, Keith et al., 2016; Momtaz et al., 2021).

In previous work, we compared children with/without AMB by evaluating time-frequency responses (i.e., neural oscillations) from multichannel EEG (Momtaz, Moncrieff et al., 2021). We showed that children with AMB had unusually large β/γ brain rhythms in response to relatively slow, click-train stimuli, suggesting a hyper-synchronization in their “neural entrainment” to complex, non-speech sounds. Entrainment is defined as the brain’s inherent ability to temporally synchronize its activity with exogenous rhythmic stimuli (Obleser & Kayser, 2019). AMB’s larger responses were accompanied by an imbalance in functional connectivity between hemispheres characterized by poor neural transmission from right to left hemisphere despite this group’s abnormally large right ear advantage behaviorally. Our previous findings led us to infer that behavioral asymmetries in children with AMB might be due to a lack of appropriate sensory processing via reduced inhibition, poorer cross-talk between auditory cortices, and especially poorer neural entrainment during even passive listening (Momtaz, Moncrieff et al., 2021). We further speculated that the inability of AMB listeners to properly entrain might result in less flexibility in how the brain adapts to changes in the sound environment, thereby rendering difficulties in extracting (or suppressing) important acoustic features needed for perception. Still, our stimulus design was limited to only a single, relatively slow click-train stimulus. Here, by explicitly manipulating the rate of stimulus presentation, we formally test our previously asserted hypothesis that AMB is associated with less flexible neural entrainment to auditory stimuli.

To this end, we recorded multichannel EEGs in children diagnosed with AMB and their age-matched peers in response to rapid non-speech stimuli. We measured neural oscillatory activity extracted from the left and right auditory cortex to assess auditory entrainment and spectrotemporal details of the EEG. Varying the rate of stimulus presentation (slow vs. fast) allowed us to directly compare the flexibility in temporal processing (i.e., adaptability to fast vs. slow stimulus rates) in AMB vs. within normal limit (WNL) children. One rate was comparable to that found in normal speech (i.e., published data from Momtaz, Moncrieff et al., 2021), while the other exceeded that of typical production. Our findings show that while WNL children easily entrain to rapid auditory stimuli, children with AMB are largely insensitive to changes in rate. Our data reveal a new AMB-related deficit in how the brain temporally tags rapid sound information.

## MATERIALS & METHODS

### Participants

The sample included n=25 children (9-12 years) who were classified into two groups [within normal limits (WNL; n=11), amblyaudia (AMB; n=14)] based on their behavioral scores on dichotic listening (DL) tests (for details, see Momtaz, Moncrieff et al., 2021). Groups were similar in age (AMB: 10.1 ± 1.7 years, WNL: 10.8 ± 1.1 years, *t*_23_ = -1.48, p = 0.15) and gender (AMB: 10/4 male/female; WNL 7/4 male/female; Fisher exact test, p = 1). None had a history of neurological impairment, head injury, chronic disease, or hearing loss (≤25 dB HL screened from 500-4000 Hz; octave frequencies). They were recruited from APD evaluation clinic referrals and flyers distributed throughout the community. Participants’ parents gave written informed consent in compliance with a protocol approved by the Institutional Review Board at the University of Pittsburgh.

### Behavioral evaluation

Three DL tests were conducted at 50 dB HL to assess binaural hearing and laterality: Randomized Dichotic Digit Test (RDDT) (Moncrieff & Wilson, 2009), Dichotic Words Test (DWT) (Moncrieff, 2015), and Competing Words subtest from the SCAN-C (CW) (Keith, 1986). Details of the behavioral evaluation are reported in Momtaz, Moncrieff et al. (2021). Briefly, right and left ear scores were converted to dominant and non-dominant so that the difference in performance between ears (reflecting interaural asymmetry) remained positive. AMB is distinguished by an abnormally large interaural asymmetry and is diagnosed when at least two dichotic listening tests indicate greater than average interaural asymmetry (Moncrieff, Keith et al., 2016).

### EEG recording procedure

#### Stimuli

Neural responses were elicited by click trains presented at two different rates. Individual clicks were 385 μ s biphasic pulses. Stimuli were presented monaurally (passive listening) at 70 dB nHL via ER-3A insert earphones. Two different presentation rates (i.e., interstimulus intervals) were used: slow (8.5/sec) and fast (14.9/sec). 1000 sweeps were collected per condition.

#### EEG recording

Data recording and analysis were identical to our previous study on AMB and neural oscillations (Momtaz, Moncrieff et al., 2021). Briefly, EEGs were recorded from 64 electrodes at 10-20 scalp locations (Oostenveld & Praamstra, 2001). Electrode impedances were <5 kΩ. EEGs were digitized using Neuroscan Synamp^2^ amplifiers at 10 kHz. Data were re-referenced to the common average offline for analysis. Continuous EEGs were processed in BESA Research 7.0 (BESA, GmbH). Recordings were epoched [-10 to 56 ms] into single trials, bandpass filtered (10-2000 Hz), and baseline corrected to the pre-stimulus interval per trial. Prior to time-frequency analysis, we rejected artifactual trials exceeding ±500 µV and those with a >75 µV amplitude gradient between consecutive samples. This resulted in 877 – 1000 artifact-free trials. Critically, trial counts did not differ between groups for either left (*t*_48_=-0.32, *p*=0.74) or right (*t*_34_=0.74, *p*=0.46) ear recordings, nor for fast (t_47_=1.11, *p*=0.26) versus slow (t_47_=-0.83, *p*=0.40) stimulus rates indicating similar overall signal-to-noise ratio.

### EEG source and time-frequency analysis

Single-trial scalp potentials were transformed into source space using BESA’s Auditory Evoked Potential (AEP) source montage (Bidelman & Momtaz, 2021). This dipole model contains regional sources in bilateral AC [Talairach coordinates (*x,y,z*; in mm): *left* = (−37, -18, 17) and *right* = (37, -18, 17)]. We extracted and averaged the time courses from the radial and tangential dipoles as these orientations capture the majority of variance describing the auditory cortical ERPs (Picton et al., 1999). This approach reduced the 64-channel data to two source dipole channels localizing current activity in left and right AC (Momtaz, Moncrieff et al., 2021). Single-trial source activity was then submitted to time-frequency analysis (TFA).

The TFA transformation was computed using a sliding window analysis on each epoch (complex demodulation; Papp & Ktonas, 1977) in 20 ms/2.5 Hz resolution step sizes (10-80 Hz bandwidth). We then computed inter-trial phase-locking (ITPL) (Lachaux et al., 1999) at each time-frequency point across single trials (Momtaz, Moncrieff et al., 2021). ITPL maps reflect the change in neural synchronization (0=random noise; 1=perfect phase-locking) relative to baseline (−10 to 0 ms) (Bidelman, 2015). Note that ITPL is invariant to amplitude (it depends only on trial phase consistency) rendering it impervious to amplitude scaling inaccuracies that might emerge from our use of adult head templates for source analysis (Momtaz, Moncrieff et al., 2021). Oscillation responses are most prominent to click train stimuli near the ∼33 Hz band of the EEG (Momtaz, Moncrieff et al., 2021). Hence, we extracted the time course of the high-β/low-γ frequency band (33 Hz) from each ITPL spectrogram. We then measured the peak ITPL strength and latency from each band time course response to quantify group effects per hemispheric source, ear, and rate of presentation.

### Statistical analysis

We used 2×2×2×2 random effects rank-based (robust) ANOVAs (R ® 4.0.3, R Foundation for Statistical Computing; Vienna, Austria) and the *lme4* (Bates et al., 2015) and *robustlmm* (Koller, 2016) R packages (R Core Team, 2020) to assess latency and ITPL strength differences in β-band responses. Note this package reports omnibus ANOVA results as *t*-(rather than *F*-) values. Fixed factors included group (2 levels: WNL, AMB ear (2 levels: LE, RE), hemispheres (2 levels: LH, RH), and rate of presentation (2 levels: fast, slow); subjects served as a random effect. The dependent variables were minimally truncated and skewed so we elected to use a robust approach to account for the distribution of these variables. Backward model selection was used to arrive at the most parsimonious model. For example, if the highest order interaction term was significant, all lower-ordered interaction terms and main effects were retained in the model. If the highest-order interaction term was insignificant, it was removed, and the next highest ordered interaction term(s) were then considered. To examine significant interactions, we stratified by the different covariates within the interaction term. The significance level was set at α= 0.05.

We used correlations (Spearman’s-*rho*) to evaluate relationships between neural oscillations (i.e., slow vs fast entrainment) and behavior (i.e., dichotic listening scores). For these analyses, a laterality index for the neural measures was computed as the *difference* in peak ITPL between ears (i.e., laterality = ITPL_RE_ -ITPL_LE_) (Jerger & Martin, 2004; Momtaz, Moncrieff et al., 2021). LH and RH responses were averaged given the lack of hemisphere effect in the omnibus ANOVAs. Neural laterality was then regressed against listeners’ three different ear advantage scores (per RDDT, DWT, and CW test), computed as the difference in behavioral performance between their dominant and non-dominant ear.

## RESULTS

A detailed analysis of the behavioral data is reported in Momtaz, Moncrieff et al. (2021). Here, we focus on new rate effects in neural entrainment in children with AMB.

### EEG time-frequency data

**Figure 1** shows ITPL spectral maps across rate, group, and hemispheres. In our backward model selection, neither the main nor interaction effects were associated with latency. However, ITPL in the β frequency band was strongly modulated by stimulus rate, ear of presentation, hemisphere, and group. Our final model using ranked based (robust) ANOVA on ITPL was: *ITPL*_*amp*_ *∼ rate + group + ear + hemi + rate*group + ear*hemi + group*hemi + (1*|*subject)*. This model revealed significant two-way interactions on neural oscillation strength including rate x group [t_169_ = -3.76, p = 0.0002], group x hemisphere [t_169_= 3.27, p = 0.001], and ear x hemisphere [t_169_= -1.99, p = 0.04] (**Fig. 2**). No other higher-order interaction terms were significant. To understand the components of these interactions, we stratified for each covariate in each two-way term.

**Figure 1:**
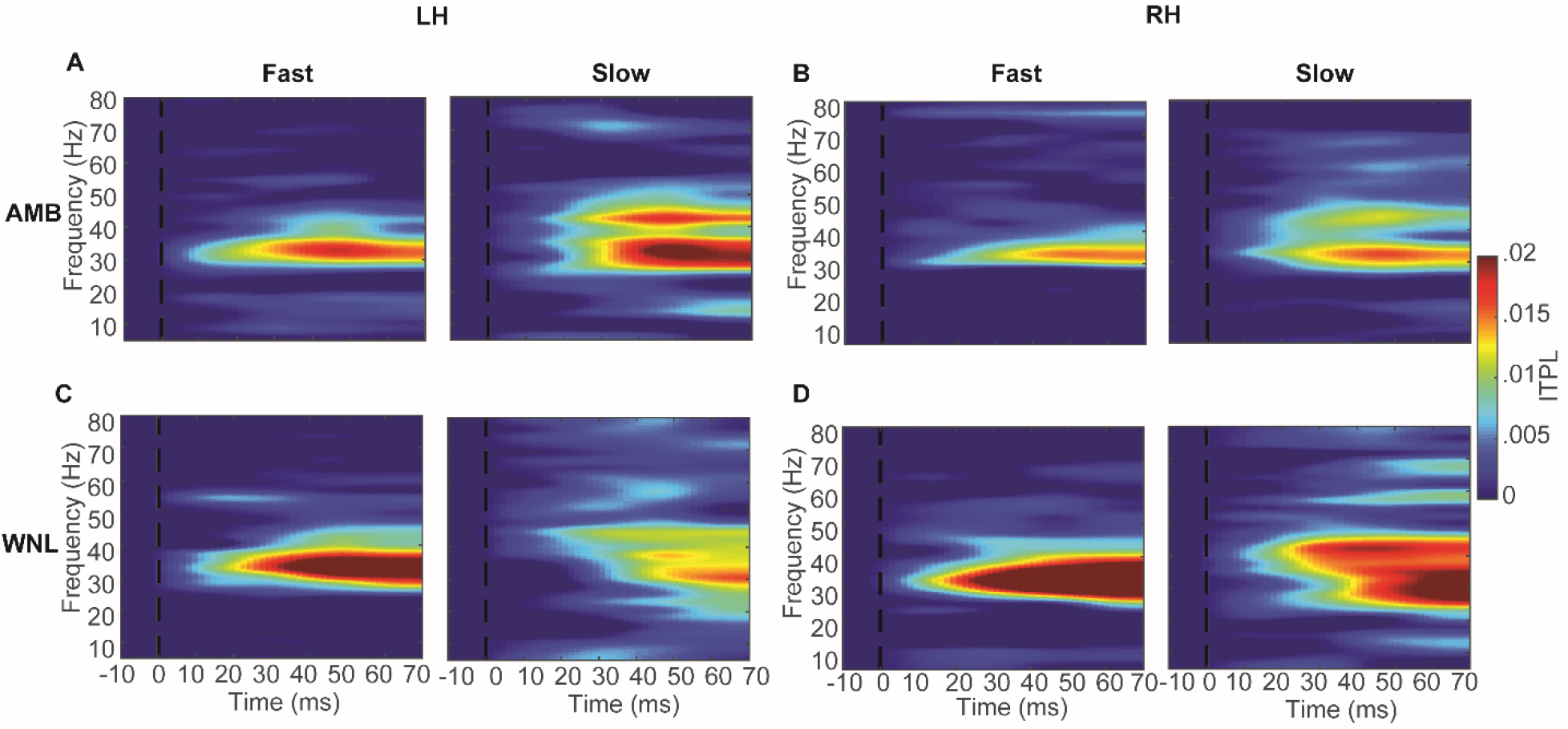
ITPL spectrograms for (**A, C**) left hemisphere and (**B, D)** right hemisphere per group and stimulus rate. Strong neural synchrony of ITPL maps is demonstrated between 30-40 Hz. AMB, amblyaudia; WNL, within normal limits; LH/RH, left/right hemisphere; ITPL, inter-trial phase-locking.

**Figure 2:**
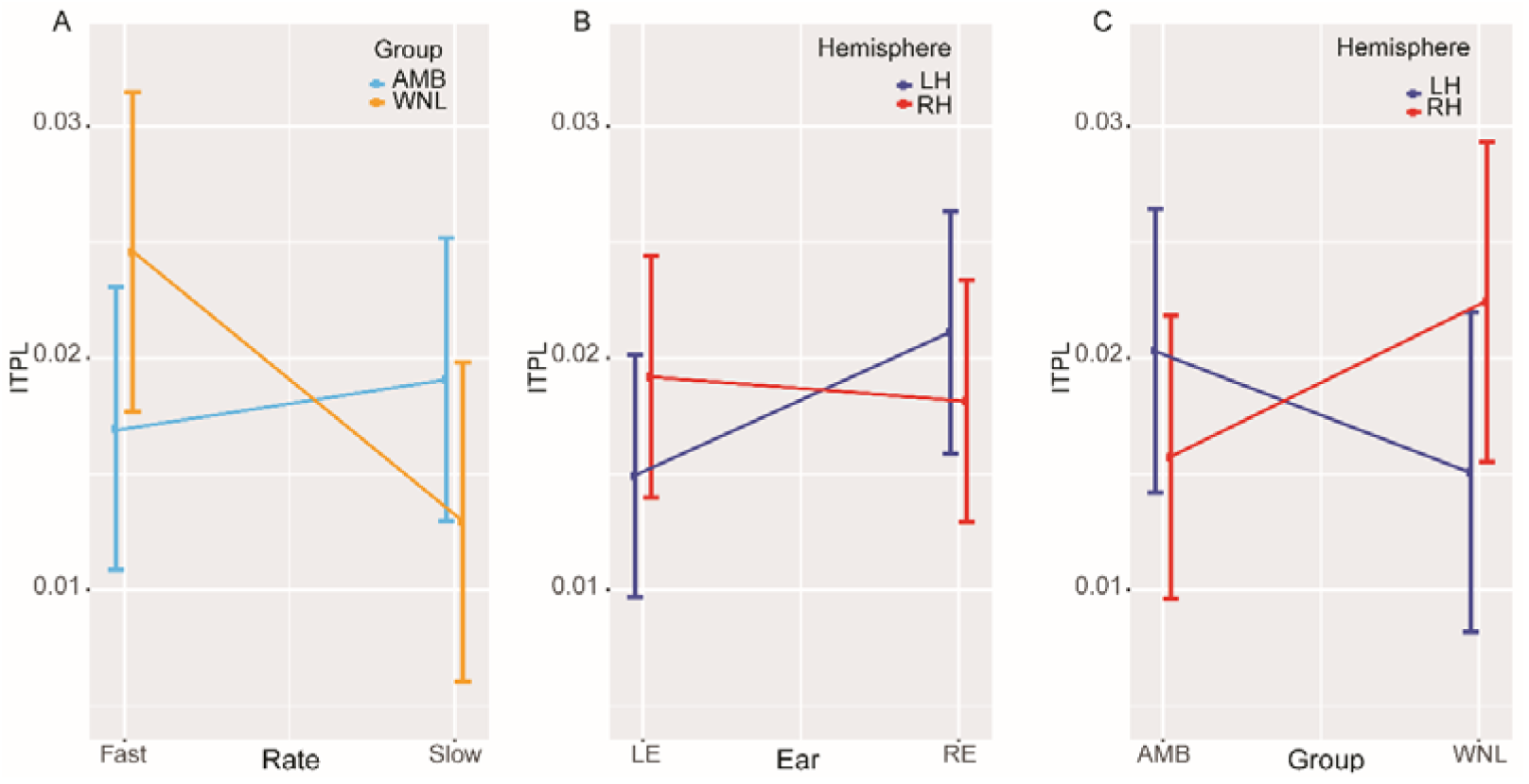
Neural oscillation strength differentially varies between groups according to stimulus rate, hemisphere, and ear. **(A)** rate*group interaction. WNLs showed decreased ITPL strength at a slow vs. fast rate. In stark contrast, no rate effects were observed in AMBs. (**B**) ear*hemisphere interaction. LH ITPL was stronger for RE vs. LE presentation for both groups. No ear differences were observed for RH responses. (**C**) group*hemisphere interaction. This interaction is stratified by ear and rate. Overall, AMB showed increased ITPL in the LH whereas WNL showed increased ITPL in the RH. error bars = ±0.95 CT.

Focusing first on **rate** stratified by group, we found stronger ITPL strength in the fast compared to the slow rate for the WNL group (p < 0.0001) but not for the AMB group (p = 0.3819). This suggests that regardless of ear and hemisphere, AMBs showed less flexibility in neural entrainment to changes in stimulus rate.

Focusing on **ear** stratified by hemisphere, we found stronger ITPL strength for RE compared to LE presentation in LH responses (p = 0.0159). No ear effect was observed in RH (p = 0.6821). These findings suggest neural responses in LH were overall stronger for right vs. left ear presentation regardless of stimulus rate and group.

We next focused on **group** stratified by both rate and hemisphere. Group differences were observed in LH at the slow rate and in RH at the fast rate. Compared to the WNL group, AMBs had stronger ITPL for slow/LH responses (p = 0.0156) but weaker ITPL for fast/RH (p = 0.0067). The group effect was not significant for the fast rate in LH (p = 0.7433) or the slow rate in RH (p = 0.9727). These findings suggest a differential pattern of neural entrainment in AMBs that varies between hemispheres and rate of stimulus presentation.

Finally, focusing on **hemisphere** stratified by group and ear, we found that RH demonstrated stronger ITPL strength among WNL for LE (p = 0.0008) compared to LH after holding rate constant. Moreover, ITPL strength was weaker among AMB for RE (p = 0.0066) compared to LH after holding rate constant. This comparison was not statistically significant among AMB in LE or among WNL in RE.

### Brain-behavior correlations

The correspondence between neural ear laterality (at slow and fast rates) and all three behavioral ear advantage scores was evaluated using correlational analysis. We previously found that the degree of ear asymmetry in neural oscillation strength to slow (8.5/sec) rate auditory stimuli were associated with behavioral performance on the DW test (Momtaz et al., 2021). However, for the fast rate (14.9/sec) stimuli in the current study, we did not find a relationship between the degree of ear asymmetry in neural oscillation strength and any of the three dichotic listening tests (all *p*s > 0.199). These results suggest that the (moderate) link between neural oscillation strength and behavioral dichotic listening (cf. Momtaz et al., 2021) is perhaps restricted to slower rate stimuli.

## DISCUSSION

Extending our prior work on the brain basis of dichotic listening deficits, we show stark differences in phase-locked neural oscillations among AMB children that depend critically on the speed of auditory stimuli. In contrast to WNL children whose neural entrainment was sensitive to rate, AMB children showed responses that were largely insensitive to rate manipulations. This resistance to rate was seen regardless of the ear of presentation and in both cortical hemispheres. Our data imply that AMB might be characterized by a varying capacity in how the brain temporally tags rapid auditory stimuli. Thus, in addition to a reduced capacity integrating information between the ears (Momtaz, Moncrieff et al., 2021), we identify a new functional characterization of AMB in the form of less flexibility (more rigidity) in how the auditory system entrains to external sounds.

### Neural entrainment differs based on the ear of presentation and hemispheres regardless of rate and group

Our results (Fig. 2B) show that right ear stimulus presentation produced stronger neural entrainment in the LH regardless of group, whereas no ear-effect was observed in RH responses. This pattern is expected given the crossed nature of the auditory neuroanatomy which leads to the typical right ear advantage (REA) and dominance in the contralateral pathway from RE to LH. Larger neural responses for the dominant contralateral auditory pathway (Jerger & Martin, 2004) regardless of stimulus properties confirm the advantage of the contralateral over the ipsilateral pathway as posited by the structural model of auditory processing (Kimura, 1967). The REA results in an interaural asymmetry of the contralateral auditory pathway that is biased to the right ear in 75-80% of right-handed and 60% of left-handed individuals, respectively (Kimura, 1967). Indeed, the majority (∼80%) of our AMB listeners were RE dominant when assessed by dichotic listening tests composed of *linguistic* materials. The fact that we find a similar REA for click entrainment suggests the REA may occur irrespective of stimulus nature (i.e., for both linguistic and non-linguistic stimuli). This finding gives credence to a “low-level deficit” account of AMB (Momtaz, Moncrieff et al., 2021). Here, we assume that the LH is more sensitive to the inputs of the dominant right ear and hence produces a larger response when the stimuli are presented to the RE as opposed to the LE. Whereas the RH mainly receives the input from the nondominant ear and hence would not be as flexible to the inputs of different ears.

### Neural entrainment in AMB is rate insensitive

Prior work shows phase-locked β/γ oscillations differ among AMB and WNL children in response to slower (8.5/sec) click-train stimuli (Momtaz et al., 2021). In particular, AMB children showed unusually large responses compared to their WNL peers. Extending those findings, new data here with faster auditory stimulation reveal a fundamentally different pattern in neural phase-locking between AMB and WNL children that is rate-dependent. Whereas control children showed changes in responses across rates (i.e., stronger ITPL strength at fast vs. slow click rates), ITPL strength was surprisingly invariant in children with AMB. Though or data reveal a rate (in)sensitivity in AMB, future studies could explore this further by evaluating ITPL with rates slower or faster than those used here to map a rate sensitivity profile (i.e., input/output function).

Differences in the resonant frequency of rhythmic entrainment (Baltus & Herrmann, 2016) could explain these group differences in the ability to entrain to rapid sounds. The precise interplay of neural excitation/inhibition can generate oscillations at a γ-band frequency that is dependent on stimulus parameters such as rate (Baltus & Herrmann, 2016). Therefore, higher stimulation rates may drive activation and boost γ oscillations, which could explain the increment in γ responses for fast vs. slow rate we find in WNLs. On the other hand, the lower ITPL strength in AMBs at faster rates could be a neurological correlate of the hallmark temporal processing deficits widely observed in APDs (Bellis, 2011). Alternatively, weaker γ responses might also be attributed to poorer perceptual-cognitive processes for rapid stimuli (Başar-Eroglu et al., 1996), which is also consistent with the dichotic and other listening difficulties observed in AMB (Momtaz, Moncrieff et al., 2021). Less robust entrainment in AMBs at faster rates also implies poorer temporal resolution in auditory processing that is perhaps analogous to a deficient pace-maker or clock that impairs brain functionality (Buzsaki, 2006). Interestingly, optimizing stimulus presentation rates shows promise in improving aspects of auditory processing (Baltus & Herrmann, 2016). Therefore, tuning external stimulus delivery to the preferred (characteristic) internal entrainment clock (which might be less flexible in AMB) might help increase the coupling between acoustic features and brain activity, ultimately leading to better and perhaps less burdensome auditory processing (e.g., Merzenich, Jenkins et al., 1996).

Across languages, the speed of conversational speech (i.e., syllable rate) unfolds at a near-universal rate between 2-8 Hz (Poeppel & Assaneo, 2020). Auditory cortical activity (Giraud et al., 2000), psychophysical performance (Viemeister, 1979), and speech comprehension (Versfeld & Dreschler, 2002) decline rapidly for modulations outside this range. Thus, both speech acoustics and auditory perception are bound by a common, fundamental upper limit of ∼8-10 Hz. Non-speech aside, our fast (14.9/sec) and slow (8.5/sec) rate stimuli might therefore be described as straddling this critical acoustic-perceptual boundary that constrains auditory temporal processing. In this regard, it is tempting to suggest that the AMB group’s stronger responses at slower rates (Fig. 2A here; Momtaz et al., 2021) might reflect the fact these stimuli are paced at the normal speech-like rate listeners are exposed to in their everyday environment. In contrast, they show inflexibility to entrain to higher (non-speech) rates where their normally developing peers show robust phase-locking. These findings parallel other electrophysiological studies showing listeners with more adept hearing skills (cf. WNL in this study) better track not only acoustic periodicities that are among their regular experiences but also do so for more complex signals that extend beyond those found in their everyday language experience (Bidelman et al., 2011). Conceivably, the relative breakdown of neural entrainment in the AMB group at higher rates might reflect the fact those sounds are faster than what is observed in everyday speech rhythms (Poeppel & Assaneo, 2020). Future studies could test this hypothesis by parametrically varying, for example, time-compressed speech.

Our stimuli were also limited to periodic signals. It is conceivable that individuals with AMB might also have difficulties nimbly switching between periodic and aperiodic acoustic events, as is characteristic of speech. Nevertheless, it should be emphasized these limits were observed using passively evoked, non-speech (repetitive click) stimuli. Consequently, we infer that temporal entrainment deficits in AMB are not restricted to speech but instead, likely reflect domain-general auditory deficits.

## Acknowledgments

We thank Drs. Karen Bell and Caitlin Price for comments on earlier versions of this manuscript. Requests for data and materials should be directed to G.M.B. This work was supported by the National Institute on Deafness and Other Communication Disorders of the NIH under award number R01DC016267 (G.M.B.), the U.S. Department of Education under award number H325K100325 (D.M.), and the Lions Hearing Research Foundation (D.M.).

## References

Baltus A. & Herrmann C.S. 2016. The importance of individual frequencies of endogenous brain oscillations for auditory cognition - A short review. Brain Res., 1640, 243–250.

Başar-Eroglu C., Strüber D., Schürmann M., Stadler M. & Başar E. 1996. Gamma-band responses in the brain: a short review of psychophysiological correlates and functional significance. International journal of psychophysiology, 24, 101–112.

Bates D., Maechler M., Bolker B. & Walker S. 2015. Fitting Linear Mixed-Effects Models Using lme4: Journal of Statistical Software, pp. 1–48.

Bellis T.J. 2011. Assessment and management of central auditory processing disorders in the educational setting: From science to practice: Plural Publishing.

Bidelman G.M. 2015. Induced neural beta oscillations predict categorical speech perception abilities. Brain Lang., 141, 62–69.

Bidelman G.M., Gandour J.T. & Krishnan A. 2011. Cross-domain effects of music and language experience on the representation of pitch in the human auditory brainstem. J. Cogn. Neurosci., 23, 425–434.

Bidelman G.M. & Momtaz S. 2021. Subcortical rather than cortical sources of the frequency-following response (FFR) relate to speech-in-noise perception in normal-hearing listeners. Neurosci. Lett., 746, 135664.

Buzsaki G. 2006. Rhythms of the Brain: Oxford University Press.

Dawes P., Sirimanna T., Burton M., Vanniasegaram I., Tweedy F., et al. 2009. Temporal auditory and visual motion processing of children diagnosed with auditory processing disorder and dyslexia. Ear and hearing, 30, 675–686.

Giraud A.L., Lorenzi C., Ashburner J., Wable J., Johnsrude I., et al. 2000. Representation of the temporal envelope of sounds in the human brain. J. Neurophysiol., 84, 1588–1598.

Grondin S. 2010. Timing and time perception: a review of recent behavioral and neuroscience findings and theoretical directions. Attention, Perception, & Psychophysics, 72, 561–582.

Jerger J. & Martin J. 2004. Hemispheric asymmetry of the right ear advantage in dichotic listening. Hear Res, 198, 125–136.

Keith R.W. 1986. SCAN: A Screening Test for Auditory Processing Disorders. San Antonio: Psychological Corporation.

Kimura D. 1967. Functional asymmetry of the brain in dichotic listening. Cortex, 3, 163–178.

Koller M. 2016. robustlmm: an R package for robust estimation of linear mixed-effects models. Journal of statistical software, 75, 1–24.

Lachaux J.P., Rodriguez E., Martinerie J. & Varela F.J. 1999. Measuring phase synchrony in brain signals. Hum. Brain Mapp., 8, 194–208.

Luthra S. 2021. The Role of the Right Hemisphere in Processing Phonetic Variability Between Talkers. Neurobiology of Language, 2, 138–151.

McFarland D. & Cacace A. 2009. Models of central auditory processing abilities and disorders. Controversies in central auditory processing disorder, 93–108.

Merzenich M.M., Jenkins W.M., Johnston P., Schreiner C., Miller S.L., et al. 1996. Temporal processing deficits of language-learning impaired children ameliorated by training. Science, 271, 77–81.

Momtaz S., Moncrieff D. & Bidelman G.M. 2021. Dichotic listening deficits in amblyaudia are characterized by aberrant neural oscillations in auditory cortex. Clinical Neurophysiology.

Moncrieff D. 2015. Age-and gender-specific normative information from children assessed with a dichotic words test. J. Am. Acad. Audiol., 26, 632–644.

Moncrieff D., Keith W., Abramson M. & Swann A. 2016. Diagnosis of amblyaudia in children referred for auditory processing assessment. Int. J. Audiol., 55, 333–345.

Moncrieff D. & Wilson R.H. 2009. Recognition of randomly presented one-, two-, and three-pair dichotic digits by children and young adults. J Am Acad Audiol, 20, 58–70.

Moore D.R. 2006. Auditory processing disorder (APD): Definition, diagnosis, neural basis, and intervention. Audiol. Med., 4, 4–11.

Obleser J. & Kayser C. 2019. Neural entrainment and attentional selection in the listening brain. Trends in cognitive sciences, 23, 913–926.

Oostenveld R. & Praamstra P. 2001. The five percent electrode system for high-resolution EEG and ERP measurements. Clin. Neurophysiol., 112, 713–719.

Papp N. & Ktonas P. 1977. Critical evaluation of complex demodulation techniques for the quantification of bioelectrical activity. Biomed. Sci. Instrum., 13, 135–145.

Picton T. 2013. Hearing in time: evoked potential studies of temporal processing. Ear and hearing, 34, 385–401.

Picton T.W., Alain C., Woods D.L., John M.S., Scherg M., et al. 1999. Intracerebral sources of human auditory-evoked potentials. Audiol. Neurootol., 4, 64–79.

Poeppel D. & Assaneo M.F. 2020. Speech rhythms and their neural foundations. Nat. Rev. Neurosci., 21, 322–334.

R Core Team 2020. R: A language and environment for statistical computing. R Foundation for Statistical Computing. Vienna, Austria.

Tallal P. 1980. Auditory temporal perception, phonics, and reading disabilities in children. Brain and language, 9, 182–198.

Toplak M.E., Dockstader C. & Tannock R. 2006. Temporal information processing in ADHD: findings to date and new methods. J. Neurosci. Methods, 151, 15–29.

Versfeld N.J. & Dreschler W.A. 2002. The relationship between the intelligibility of time-compressed speech and speech in noise in young and elderly listeners. J. Acoust. Soc. Am., 111, 401–408.

Viemeister N.F. 1979. Temporal modulation transfer functions based upon modulation thresholds. J. Acoust. Soc. Am., 66, 1364–1380.

